# A role for the ATP-dependent DNA ligase Lig E of *Neisseria gonorrhoeae* in biofilm formation

**DOI:** 10.1101/2023.09.19.558522

**Authors:** Jolyn Pan, Joanna Hicks, Adele Williamson

## Abstract

The ATP-dependent DNA ligase Lig E is present as an accessory DNA ligase in numerous proteobacterial genomes, including many disease-causing species. Here we have constructed a genomic Lig E knock-out in the obligate human pathogen *Neisseria gonorrhoeae* and characterised its growth and infection characteristics. This demonstrates that *N. gonorrhoeae* Lig E is a non-essential gene and its deletion does not cause defects in replication or survival of DNA-damaging stressors. Knock-out strains were partially defective in biofilm formation on an artificial surface as well as adhesion to epithelial cells which coupled with the predicted extracellular/ periplasmic location of Lig E indicates a role in extracellular DNA joining. In addition to *in vivo* characterisation, we have recombinantly expressed and assayed *N. gonorrhoeae* Lig E and determined the crystal structure of the enzyme-adenylate engaged with DNA substrate in an open non-catalytic conformation, providing insight into the binding dynamics of these minimal DNA ligases.

## INTRODUCTION

The DNA ligase Lig-E is the most recently-delineated form of the diverse bacterial ATP-dependent DNA ligases (b-ADLs) which are found in the genomes of many bacterial species in addition to their replicative NAD-dependent DNA ligase. The b-ADLs characterised to date are auxiliary enzymes which join DNA breaks as part of stationary phase DNA-repair pathways. These structurally diverse enzymes are categorised by both the composition of their appending domains, some of which have autonomous catalytic functions and their operonic arrangement with other pathway enzymes. The best-studied b-ADLs Lig C and Lig D function in stationary-phase base excision repair and non-homologous end joining respectively and are found adjacent to other genes that carry out earlier nucleolytic, polymerase or DNA end-binding steps ^1-3^. Likewise, the co-localization of the Lig B class of b-ADLs with a conserved Lhr-helicase and pair of nucleases implies a role in genomic DNA repair ^4,5^.

In contrast to these b-ADLs, Lig E does not exhibit syntetic conservation with any repair-associated genes and its biological function remains undefined ^6,7^. Lig E is distinguished by having a minimal structure lacking typical globular DNA binding domains or loop regions. Despite this, *in vitro* characterisation of recombinant Lig E enzymes from a range of bacteria have demonstrated that they are fully-functional ATP-dependent ligases with preferential activity on singly-nicked DNA and some, albeit weaker, activity on cohesive ends ^8-11^. Structures of Lig E bound to adenylated nicked DNA or as the enzyme-adenylate without DNA show that it engages this substrate through highly conserved basic residues in the oligonucleotide binding (OB) domain and the inter-domain linker ^9,10^. The most intriguing structural aspect of Lig E however, is the N-terminal leader sequence which is predicted to direct its localisation to the periplasm. Removal of this sequence increases both the stability and activity of the enzyme, and both Lig E structures as well as most *in vitro* characterisation has been undertaken on the mature leader-less form ^11^.

Examination of the Phylogenetic distribution of Lig E indicates it is widespread among, but restricted to proteobacteria and is typically the only b-ADL found in a particular species’ genome ^7^. Notably, Lig E has been annotated in genomes of several naturally competent, biofilm-forming pathogenic bacteria, several of which have acquired antibiotic resistance traits and present potential multi drug resistance threats ^6^. An example is the obligate human pathogen *Neisseria gonorrhoeae* which colonises and infects mucosal cells of the male and female reproductive tracts and is the causative agent of the sexually transmitted infection (STI), gonorrhoea. One of the most concerning features of *N. gonorrhoeae* is its ability to gain antibiotic resistance due to its natural competence. Its propensity to take up conspecific DNA is enhanced 20 – 100-fold by the presence of a 10 bp DNA uptake sequence (DUS) which is common in the gonococcal genome and allows the bacterium to differentiate species-specific DNA from other DNA in the environment ^12,13^. Gonorrhoea infection may cause urethritis in men, cervicitis or pelvic inflammatory disease in women and neonatal conjunctivitis if contracted during birth; however, a high proportion of infections are asymptomatic (approximately 50% in females and 40% in males ^14^) leading to its undetected spread and delayed treatment ^15^. *N. gonorrhoeae* readily forms a biofilm on epithelial cells during infection which likely contributes to this ability to evade the host immune system through the help of a heterogeneous physical barrier or the potential subsequent induction of oxidative stress defence mechanisms, and additionally can modulate the spread of antibiotic resistance by horizontal gene transfer ^16-19^. Extracellular DNA (eDNA) comprises a large fraction of gonococcal biofilm which is produced by active DNA secretion as well as contribution of genomic DNA from lysed gonococcal cells ^20-22^.

Despite extensive *in vitro* characterisation of Lig E from a range of pathogenic and environmental bacteria, the biological function of Lig E remains unknown. However, the probable extracellular location of Lig E, coupled with its presence in the genomes of bacteria known to be competent for eDNA uptake and which can form eDNA-rich biofilms suggest its involvement of this DNA ligase in one or both of these processes. Here we report the first *in vivo* study of the function of Lig E via generation of a knock-out in *N. gonorrhoeae*.

Although no impact was observed on planktonic growth rates, the deletion mutant was partially defective in biofilm formation and adhesion to epithelial cells as well as exhibiting aberrant growth rates in the presence of DNA-damaging antibiotics. In addition, we have characterised recombinantly-produced *N. gonorrhoeae* Lig E *in vitro* and determined the structure of the enzyme-adenylate in an open DNA-engaged conformation.

## METHODS

### *N. GONORRHOEAE* STRAINS AND CULTIVATION

All *N. gonorrhoeae* used in this study were generated from the MS11 strain. Gonococci were grown at 37 °C and 5 % CO_2_ either on gonococcal base (GCB) agar (Difco) or in gonococcal base liquid (GCBL) media (1.5 % (w/v) proteose peptone #3, 0.4 % (w/v) K_2_HPO_4_, 0.1 % (w/v) KH_2_PO_4_, 0.1 % (w/v) NaCl), both supplemented with 1 % *Kellogg’s* supplement (22.22 mM glucose, 0.68 mM glutamine, 0.45 mM cocarboxylase, 1.23 mM Fe(NO_3_)3) ^23^.

### *N. GONORRHOEAE* MUTANT CONSTRUCTION

To generate different variants, constructs were designed to introduce insertions and/or deletions into the MS11 genome (GenBank: CP003909.1) by homologous recombination of flanking sequences. Briefly, the *Δngo-Lig E* mutant contained a disruption of the *ngo-Lig E* gene (NGFG_01849) via a kanamycin resistance cassette, while the *ngo-Lig E-His* mutant had a 6-His-tag at the C-terminus of the intact *ngo-Lig E* gene as well as an additional kanamycin resistance cassette behind the gene. The *opaB-ngo-Lig E* mutant was generated at an intergenic site between open reading frames NGFG_RS14495 and NGFG_RS13185.

Here, a codon-optimised *ngo-Lig E* gene was used to avoid aberrant recombination with the native *ngo-Lig E* copy while still encoding the same amino acid sequence. This was inserted behind the constitutive *opaB* promoter and included a 6-His-tag at the C-terminus and a kanamycin resistance cassette for selection. All DNA constructs were ordered as gene fragments or clonal genes (Integrated DNA Technologies or Twist Biosciences).

Strains were generated via spot transformation ^23,24^. Briefly, piliated colonies were streaked through 10 ng spots of the DNA constructs on GCB agar. After a 24 hr incubation, colonies growing at the spotted locations were restreaked onto GCB plates with kanamycin (50 μg/mL) for selection. These mutants were verified via PCR and sequencing analyses using primers detailed in Supplement 1.

### GROWTH EXPERIMENTS IN LIQUID CULTURE

Pilliated gonococci from a 24 hr streak were lawned for 16 hr before resuspension in GCBL media. Suspensions with a OD_600_ of 0.05 were prepared and aliquoted into 12-well plates (1 mL per well, 3 replicates each), where each 12-well plate corresponded to one time point.

Gonococcal cells were harvested at 1.5 hr intervals by scraping cells from the bottom of the well and vortexing vigorously for 2 min. Growth was monitored by measuring the OD_600_ of the cell resuspensions before serially diluting and plating onto GCB agar. The number of colonies on the agar plates were counted after 48 hrs to obtain colony forming units (CFUs).

### H_2_0_2_ OXIDATIVE STRESS ASSAY

Pilliated gonococci from a 24 hr streak were lawned for 16 hr before resuspension in GCBL media. Suspensions with a OD_600_ of 0.05 were prepared and aliquoted into separate wells in 12-well plates. After 9 hr of growth, the gonococci were subjected to 0, 2, 5, 10, 25 or 50 mM H_2_0_2_ treatment for 20 min. Cells were then scraped from the bottom of the wells, pelleted and washed with 300 μL GCBL to remove excess hydrogen peroxide, then resuspended in 1 mL GCBL before being serially diluted and plated onto GCB agar. CFU readings were obtained by counting the number of colonies formed after 48 hr.

### UV SURVIVAL ASSAY

Pilliated gonococci from a 24 hr streak were lawned for 16 hrs before resuspension in GCBL media. Suspensions with an OD_600_ suspension of 0.6 were prepared and serially diluted before plating onto GCB agar. The agar plates were subjected to UV irradiation at 80 J for 0, 5, 7.5 and 10 min using a BLX-254 crosslinker. The plates were then incubated for 48 hrs before counting to obtain CFU readings.

### NALIDIXIC ACID TREATMENT ASSAY

Pilliated gonococci from a 24 hr streak were lawned for 16 hrs before resuspension in GCBL media. Suspensions with an OD_600_ of 0.6 were prepared and serially diluted before plating onto GCB agar with 1.25 mg/L nalidixic acid (Nal-acid). The plates were incubated for 48 hr before counting to obtain CFU readings.

### CELL INFECTION AND ADHESION ASSAYS

The ME-180 endocervical cell line was used for the host association assays and were maintained in McCoy’s 5A media (Gibco) supplemented with 10 % foetal bovine serum (FBS). ME-180 cells were seeded in 12-well plates 48 hr prior to use to achieve 80-100 % confluency on the day of the experiment. Pilliated, Opa negative (Opa^-^) gonococci from a 24 hr streak were lawned for 16 hrs before resuspension in GCBL media. The OD_600_ of the resuspensions were measured and back-calculated to give units of CFU/mL. The suspensions were then used to infect the ME-180 cells at a multiplicity of infection (MOI) of 25 in McCoy’s 5A with 10 % FBS for 6 hr. All experiments were done in triplicate.

For planktonic measurements, the supernatants were aspirated and the wells were washed three times with 1 mL GCBL. The pooled supernatant and washes were vortexed (2 min), serially diluted and plated onto GCB agar. To measure adhered cells, the remaining cell monolayers were subjected to 0.5 % saponin treatment (1 mL in GCBL) for 20 min then scrapped and vortexed vigorously for 2 min before serial dilution and plating onto GCB agar. The number of colonies on the agar were counted after 48 hours to obtain CFU readings for planktonic and adhered cells respectively. Gonococcal adherence and planktonic growth were calculated as the proportions of the total CFUs and expressed as percentages.

For invasion measurements, the media was aspirated before treatment of infected ME-180 cells with 50 μg/mL gentamicin for 1 hr. The wells were then washed three times with 1 mL GCBL before being subjected to 0.5% saponin (1 mL in GCBL) for 20 min. The cells were then scraped and vortexed vigorously for 2 min before serial dilution and plating onto GCB agar. The number of colonies on the agar were counted after 48 hr to obtain CFU readings for cells that had invaded the cervical cell monolayer. The extent of invasion was calculated as a proportion of the total number of cells.

### BIOFILM MICROTITER ASSAYS

Pilliated gonococci from a 24 hr streak were lawned for 16 hr before resuspension in GCBL media. Suspensions with an OD_600_ of 0.05 were prepared and aliquoted into separate wells in 96-well plates (100 μL per plate, 8 replicates each). After 24 hr, the wells were washed three times with sterile water before staining with 125 μL 0.8 % crystal violet for 15 min. The wells were then washed four times with sterile water before air-drying overnight. The dye was resolubilised in 125 uL 30 % acetic acid and the solubilised crystal violet solutions were transferred to a new 96-well plate. The extent of biofilm formed was determined by the absorbance at 560 nm.

### RNA EXTRACTION AND RT-QPCR

RNA was isolated from WT gonococci and the three variant strains under planktonic and biofilm conditions. Pilliated gonococci from a 24 hr streak were lawned for 16 hr before resuspension in GCBL media. Suspensions with an OD_600_ of 0.05 were prepared and aliquoted into separate wells in 12-well plates. After 24 hr, the cells were either harvested without scraping (planktonic fraction) or harvested with scraping the wells (biofilm fraction). Total RNA was isolated using the Direct-zol RNA Miniprep Kit (Zymo Research). RNA concentration and quality were measured using the DeNovix DS-11 spectrophotometer and the Denovix RNA quantification assay kit. Reverse transcription was performed on 18 ng/μL RNA to obtain cDNA using the SuperScriptIII First-Strand Synthesis System (Invitrogen).

RT-qPCR was performed on a Mic qPCR cycler using the Hot Fire Pol DNA polymerase kit (Solis Biodyne) with specific probes and primers for the Lig E gene and the 16s rRNA housekeeping gene as listed in Supplement 2. Relative quantification of gene transcription was performed using the comparative Ct method ^25^ after normalising to the 16s rRNA gene.

### SUBCELLULAR FRACTIONATION

Subcellular fractionation was performed on gonococcal cells to separate cytoplasmic, cell-membrane, periplasmic and extracellular proteins. Pilliated gonococci from a 24 hr streak were lawned for 16 hr before resuspension in GCBL media. 30 mL cultures with an OD_600_ of 0.05 were prepared with GCBL media and cultivated overnight before harvesting by centrifugation (5000 xg, 15 min) to separate the pellet and supernatant. Extracellular proteins were recovered from the supernatant by precipitation with 20 % trichloroacetic acid, incubation for 1 hr on ice and collection by centrifugation at 20,000 xg. The resultant pellet was washed with ice-cold 90% acetone three times before air drying and resuspension in 10 mM Tris (pH 8.0). The periplasmic fraction was isolated from the pelleted cells by addition of 1 mL of buffer 1 (0.2 M Tris (pH 8.0), 0.1 M EDTA, 20 % sucrose) before incubation on ice (20 min) and centrifugation (20,000 xg 15 min, 4 °C). The pellet was resuspended in 1 mL buffer B (10 mM Tris, 5 mM MgSO_4_, 0.2 % SDS, 1 % Triton X100) before incubation on ice (20 min) and centrifugation (20,000 xg, 15 min, 4 °C). The resultant supernatant was the periplasmic portion. To isolate cytoplasmic fraction proteins, the remaining pellet was treated with 1 mL Bug Buster (Sigma-Aldrich) and agitated for 20 min before centrifugation (10000 rpm, 10 min, 4 °C). The supernatant was re-centrifuged at maximum speed (1 hour) and the resultant supernatant was isolated as the cytoplasmic portion. To isolate the membrane fraction isolation, the remaining pellet was resuspended in 0.01 M Tris (pH 8), spun and the resultant pellet was isolated as the membrane portion

### HIS-TAGGED PROTEIN DETECTION

To enrich for His-tagged proteins, each subcellular fraction was incubated with pre-washed Ni Sepharose High Performance nickel resin beads (Cytiva) for 15 min. After this time, the beads were sedimented by centrifugation, washed twice with lysis buffer (50 mM Tris pH 8.0, 750 mM NaCl, 1 mM MgCl_2_, 5 % glycerol) and electrophoresed on 12 % SDS-PAGE gels.

Western blotting was performed with nitrocellulose membranes. After protein transfer, membranes were blocked for an hour with 5 % milk in Tris buffered saline-Tween 20 (TBST). The membrane was probed with 1:500 anti-His-tag mouse monoclonal (HIS.H8), sc57598 igG2b antibody (Santa Cruz Biotechnology; 10 μg/mL) overnight, and 1:1000 goat anti-mouse polyclonal IgG antibody conjugated to horseradish peroxidase ab97023 (Abcam; 1 mg/mL) for 1 hr. The membranes were incubated with the SuperSignal West Femto Maximum Sensitivity Substrate for 5 min before imaging using the iBright Imaging System.

### RECOMBINANT EXPRESSION OF NGO-LIG E

The position of the Ngo-Lig E N-terminal leader sequence was predicted using Signal P ^26^. Pre-cloned constructs encoding mature native- and C-terminally His-tagged Ngo-Lig in the pDONR221 plasmid were synthesised from Twist BioScience with codon optimization for *E. coli*. Constructs were sub-cloned into the pDEST 17 and pHMGWA vectors using the Gateway system and recombinant Ngo-Lig E was expressed and purified from BL21(DE3)pLysS at 15 °C as described for other Lig E proteins previously ^11^.

Briefly, native mature Ngo-LigE expressed from pDEST17 with an N-terminal His tag was purified with a primary immobilised metal affinity chromatography (IMAC) step on a 5 mL His trap HP column with buffer A (50 mM Tris pH 8, 750 mM NaCl, 10 mM imidazole, 5% glycerol) and eluted with buffer B (50 mM Tris pH 8, 750 mM NaCl, 500 mM imidazole, 5% glycerol). After exchange into TEV cleavage buffer C (50 mM Tris pH 8, 100 mM NaCl, 5% glycerol, 1 mM DTT) the N-terminal His-tag was cleaved overnight with TEV protease (0.1 mg/ml) and the de-tagged protein was recovered by a reverse IMAC step. A final size-exclusion chromatography (SEC) was carried out using a Hi Load 16/600 Superdex 75 column. Native mature Ngo-LigE and C-terminally tagged mature Ngo-LigE expressed with N-terminal His-MBP tags were purified in the same way, but an additional chromatographic step was included after size exclusion to separate residual His-MBP tag that had carried over after cleavage. Pooled Ngo-Lig E/ Ngo-Lig E-His were loaded onto an MBPTrap HP column in MBP binding buffer (20 mM Tris pH 7.4, 200 mM NaCl, 1 mM EDTA, 1 mM DTT) and eluted using a linear gradient of MBP elution buffer (20 mM Tris pH 7.4, 200 mM NaCL, 1 mM EDTA, 1 mM DTT, 10 mM Maltose). All proteins were evaluated as being purified to homogeneity by the appearance of a single band on SDS-PAGE.

### DNA LIGATION ASSAYS

Gel based endpoint assays were used to measure ligation activity as described previously ^6,27^. Standard assay conditions included 80 nM of fluorescently-labelled nicked substrate, 1.0 mM ATP, 10 mM MgCl_2_, 10 mM DTT, 50 mM NaCl and 50 mM Tris pH8.0. Ngo-Lig E or Ngo-Lig E-His (0.1 μM) were incubated at 25 °C for 30 min before quenching with 95 % formamide stop buffer. Products were electrophoresed on 20 % urea-PAGE gels and fluorescence was detected using the iBright imaging system and quantified using Image J ^28^. The assay was repeated with variations in the pH (Tris buffer for pH 7.1-9.0; MES buffer for pH 5.5-6.2) and amount of ATP used in the reaction buffer as well as different combinations of substrate oligonucleotides to generate different ligatable DNA breaks (Supplement 3 and Supplement 4). Incubation conditions for the different DNA substrates were 25 °C, 30 min for single nick, overhang and mismatched substrates, and 15 °C overnight for blunt ended and gapped substrates.

### CRYSTALLIZATION AND STRUCTURE DETERMINATION OF THE NGO-LIG E – DNA COMPLEX

Double-stranded nicked DNA for co-crystallization was assembled as described previously ^6,9^ using HPLC-purified oligos purchased from IDT with the sequences CAC TAT CGG AA (5’P-phosphorylated strand); ATT GCG ACC (3’OH-strand) and TTC CGA TAG TGG GGT CGC AAT (complementary strand). His-tagged Ngo-Lig E (478.7 μM) was incubated with a 1.2 molar excess of the nicked duplex DNA an additional 5 mM EDTA for 1 hr on ice prior to commencing crystallization screening. Crystals with a plate morphology were grown by hanging drop diffusion at 18 °C in 0.5 M potassium thiocyanate, 0.1 M Bis Tris Propane pH 8.0 and were mounted in cryoloops and directly flash frozen in liquid nitrogen for data collection. Diffraction data to 2.44 Å was measured at the Australian Synchrotron MX2 beamline ^29^ and integrated, scaled and merged using XDS and Aimless ^30,31^. A model of Ngo-Lig E was built using AlphaFold via the CoLab server ^32^ and processed using the Process Predicted Model utility in the Phenix suite ^33^. The processed NTase and OB domains were used as search models for molecular replacement in Phaser-MR ^34^ together with iteratively-truncated portions of double-stranded DNA from the Ame-Lig co-crystal (6gdr). The initial model was improved by iterative rounds of refinement using Phenix.refine ^35^ and manual rebuilding in COOT ^36^. Data collection and statistics are listed in Table 1 and the structure was deposited to the Protein Data Bank with the identifier 8U6X.

**Table 1.** Data collection and refinement statistics. Statistics for the highest-resolution shell are shown in parentheses.

|  | Ngo-Lig E (8U6X) |
| --- | --- |
| Wavelength | 0.9537 |
| Resolution range | 43.67 - 2.44 (2.527 - 2.44) |
| Space group | P 21 21 2 |
| Unit cell | 39.398 167.68 51.159 90 90 90 |
| Total reflections | 115570 (11213) |
| Unique reflections | 13286 (1269) |
| Multiplicity | 8.7 (8.8) |
| Completeness (%) | 99.86 (99.61) |
| Mean I/sigma(I) | 11.88 (1.64) |
| Wilson B-factor | 53.04 |
| R-merge | 0.1076 (1.08) |
| R-meas | 0.1144 (1.146) |
| R-pim | 0.03774 (0.3757) |
| CC1/2 | 0.997 (0.694) |
| CC* | 0.999 (0.905) |
| Reflections used in refinement | 13273 (1266) |
| Reflections used for R-free | 1329 (127) |
| R-work | 0.2301 (0.3423) |
| R-free | 0.2809 (0.3688) |
| CC(work) | 0.937 (0.764) |
| CC(free) | 0.891 (0.577) |
| Number of non-hydrogen atoms | 2295 |
| macromolecules | 2222 |
| ligands | 43 |
| solvent | 42 |
| Protein residues | 247 |
| RMS(bonds) | 0.003 |
| RMS(angles) | 0.51 |
| Ramachandran favored (%) | 95.92 |
| Ramachandran allowed (%) | 4.08 |
| Ramachandran outliers (%) | 0.00 |
| Rotamer outliers (%) | 2.96 |
| Clashscore | 3.21 |
| Average B-factor | 61.64 |
| macromolecules | 58.78 |
| ligands | 66.12 |
| solvent | 44.75 |

## RESULTS

### NGO-LIG E IS NOT ESSENTIAL FOR GONOCOCCAL GROWTH AND SURVIVAL

To evaluate the importance of Ngo-Lig E for *N. gonorrhoeae* viability, physiology and stress survival, we constructed a knock-out of the *ngo-Lig E* gene (*Δngo-Lig E*) by interruption of the *ngo-lig E* open reading frame with a kanamycin resistance cassette (*Kan*^*R*^) which removed a stretch of 795 nucleotides from the centre of the gene (Figure 1 A). A second construct was generated which appended a 6-His-tag to the C-terminus of native Ngo-Lig E and inserted a Kan^R^ cassette behind the *ngo-Lig E* gene (*ngo-Lig E-His*). The purpose of this was to provide an immunogenic handle on natively-produced Ngo-Lig E, and this construct additionally served as a control for the knock-out with both strains containing equivalent Kan^R^ insertions in their genomes. A third construct (*opaB-ngo-Lig E*) generated a constitutive high-expression genotype, by inserting a second copy of His-tagged *ngo-Lig E* under the control of the strong constitutive promoter *opaB* at a separate location in the genome between a PLxRFG (NGFG_RS14495) domain containing protein and a hypothetical protein (NGFG_RS13185). The resulting clones were sequenced to confirm the correct genotypes. qPCR indicated a 89-fold upregulation of *ngo-lig E* transcripts from the *opa-ngo-Lig E* strain and confirmed that the *ngo-Lig E* gene expression was eliminated from the *Δngo-Lig E* strain (Supplement 5 and Figure 2 A). Growth in liquid culture measured by both OD_600_ and CFU counts showed no significant difference between wild-type *N. gonorrhoeae* (*wt*), *Δngo-Lig E, ngo-Lig E-His* and *opa-ngo-Lig E* which suggests that *ngo-Lig E* is neither an essential gene, nor is its overexpression deleterious to cell survival in planktonic culture (Figure 1 B and C). Attempts were made to visualize Ngo-Lig E-His expression by Western blot against the 6-His tag in both *ngo-Lig E-His* and *opa-ngo-lig E* during different growth stages; however despite evidence of gene expression by qPCR, neither strain showed detectable immunologic signal by this method (Supplement 6).

**Figure 1.**
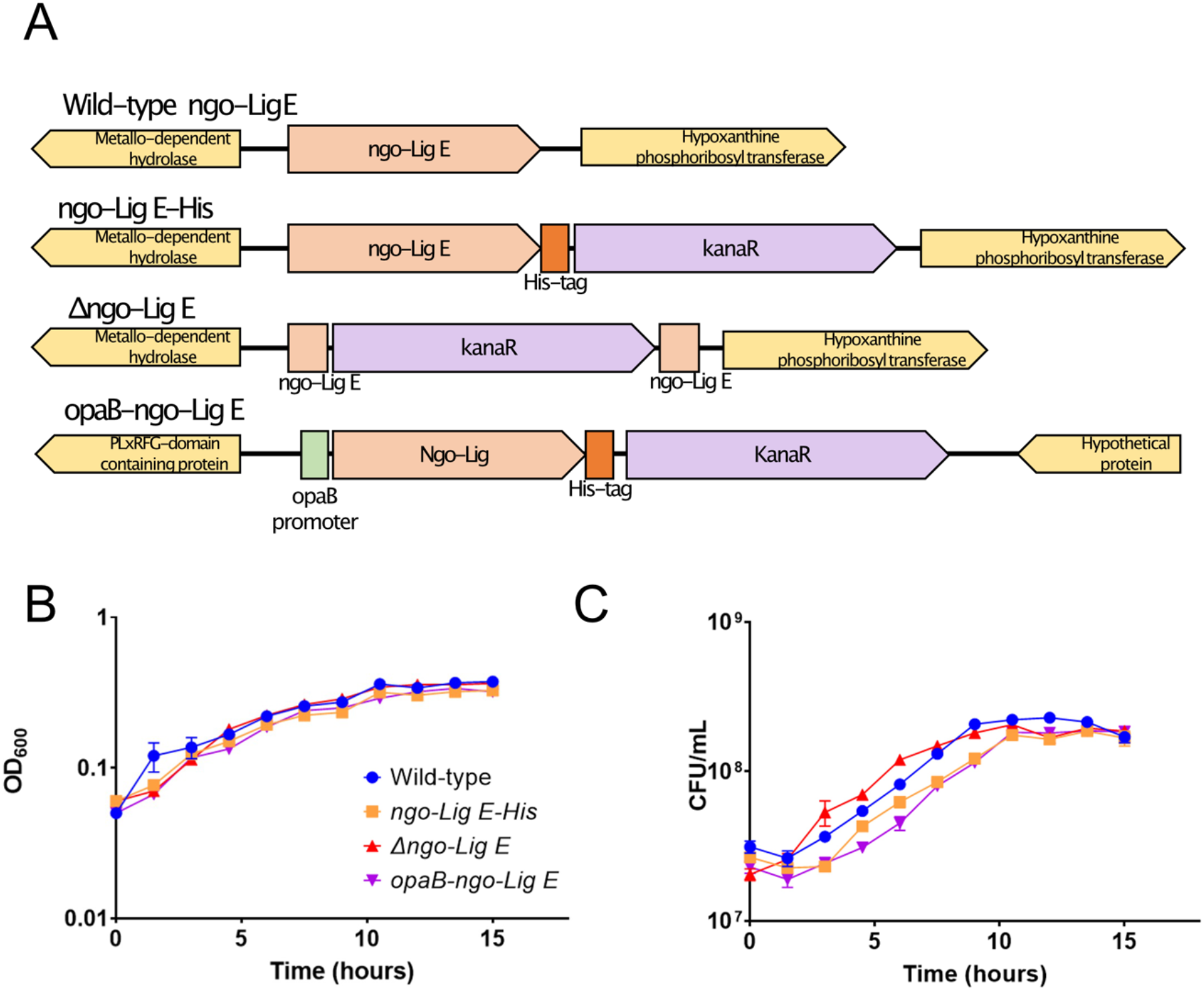
A) Schematic of the genetic constructs for His-tagged Ngo-lig E (ngo-Lig E-His), the Ngo-lig E deletion mutant (Δngo-Lig E) and constitutively upregulated Ngo-Lig E (opaB-Ngo-Lig E). B) Growth of N. gonorrhoeae variants monitored by OD_600_. C) Number of viable cells of N. gonorrhoeae variants monitored by CFU counts for each culture. Points are the mean of triplicate measurements and error bars represent the standard error of the mean.

**Figure 2.**
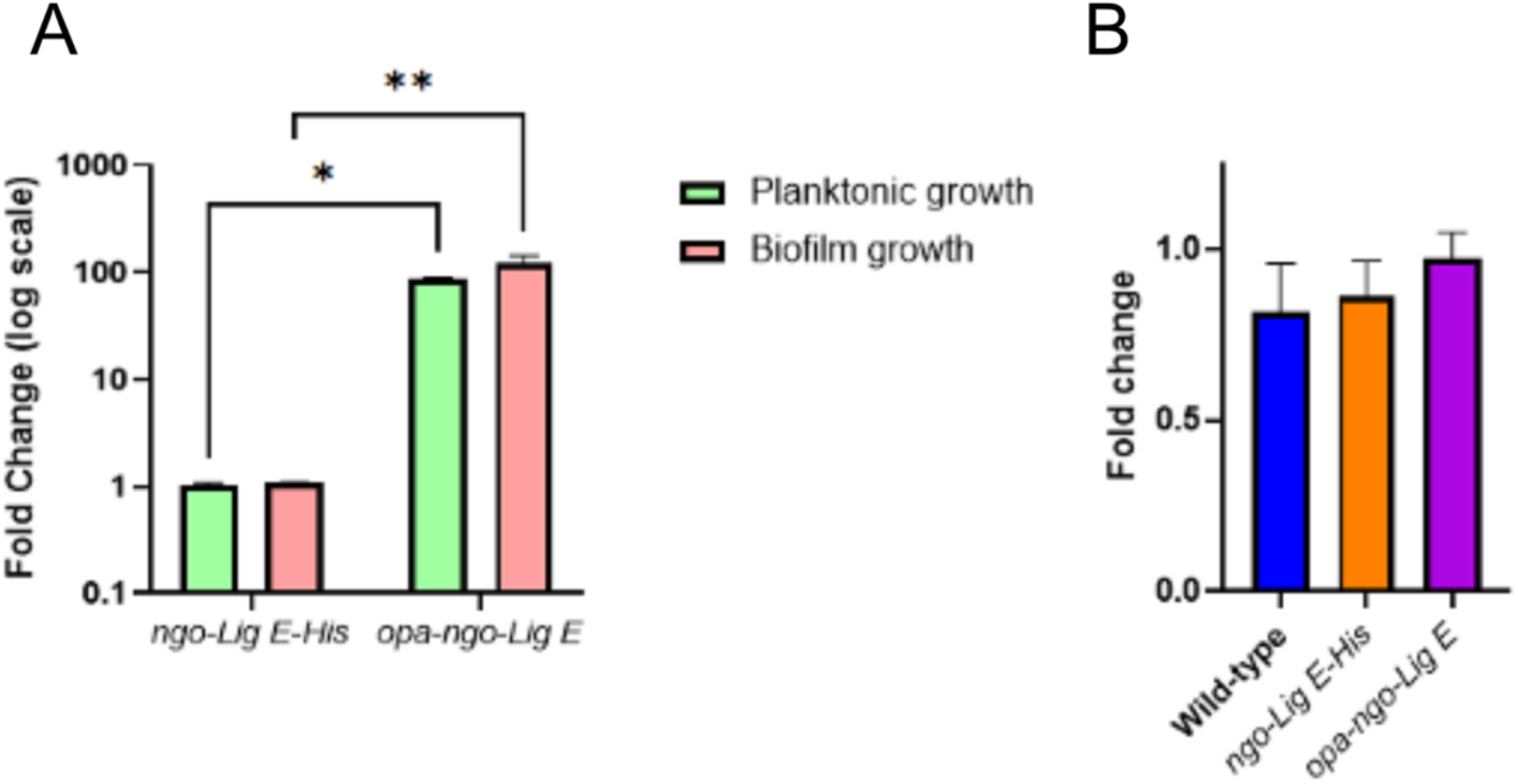
Comparisons of gene expression with normalisation to the 16S gene. A) Fold changes of ngo-Lig E-His and opa-ngo-Lig E compared to wild-type N. gonorrhoeae under either planktonic or biofilm conditions. B) Fold changes of wild type, ngo-Lig E-His, and opa-ngo-Lig E gene expression under biofilm conditions compared to planktonic conditions. Points are the mean of triplicate measurements and error bars represent the standard error of the mean. Significance values are given as * P ≤ 0.05; ** P ≤ 0.01. Comparisons which showed no significant difference (P >0.05) are not indicated.

### RESPONSE TO OXIDATIVE AND DNA-DAMAGING STRESSORS

Given the role of DNA ligases in genomic DNA repair and replication, we asked whether Ngo-Lig E plays a role in surviving DNA damage or oxidative stress in *N. gonorrhoeae* by subjecting the *Δngo-Lig E, ngo-Lig E-His* and *opa-ngo-lig E* strains and *wt* to genotoxic stressors. No significant differences in survival were seen between either the Lig E deficient *Δngo-Lig E* strain, or the *opa-ngo-lig E* over expresser when exposed to increasing concentrations of hydrogen peroxide or UV dosages (Figure 3 A and B). As H_2_O_2_ and UV are both expected to damage chromosomal DNA, the absence of higher mortality in either of these strains suggests that Ngo-Lig E does not play a significant role in repair of chromosomal DNA damage. A significant increase in survival was observed for the *Δngo-Lig E* mutant when treated with nalidixic acid, with almost twice as many cells surviving at a concentration of 1.25 mg/L relative to the *wt* (Figure 3 C). Nalidixic acid induces double-stranded breaks in chromosomal DNA, suggesting that although Ngo-Lig E may not be involved directly in intracellular DNA repair processes, it is still able to interact with chromosomal DNA.

**Figure 3.**
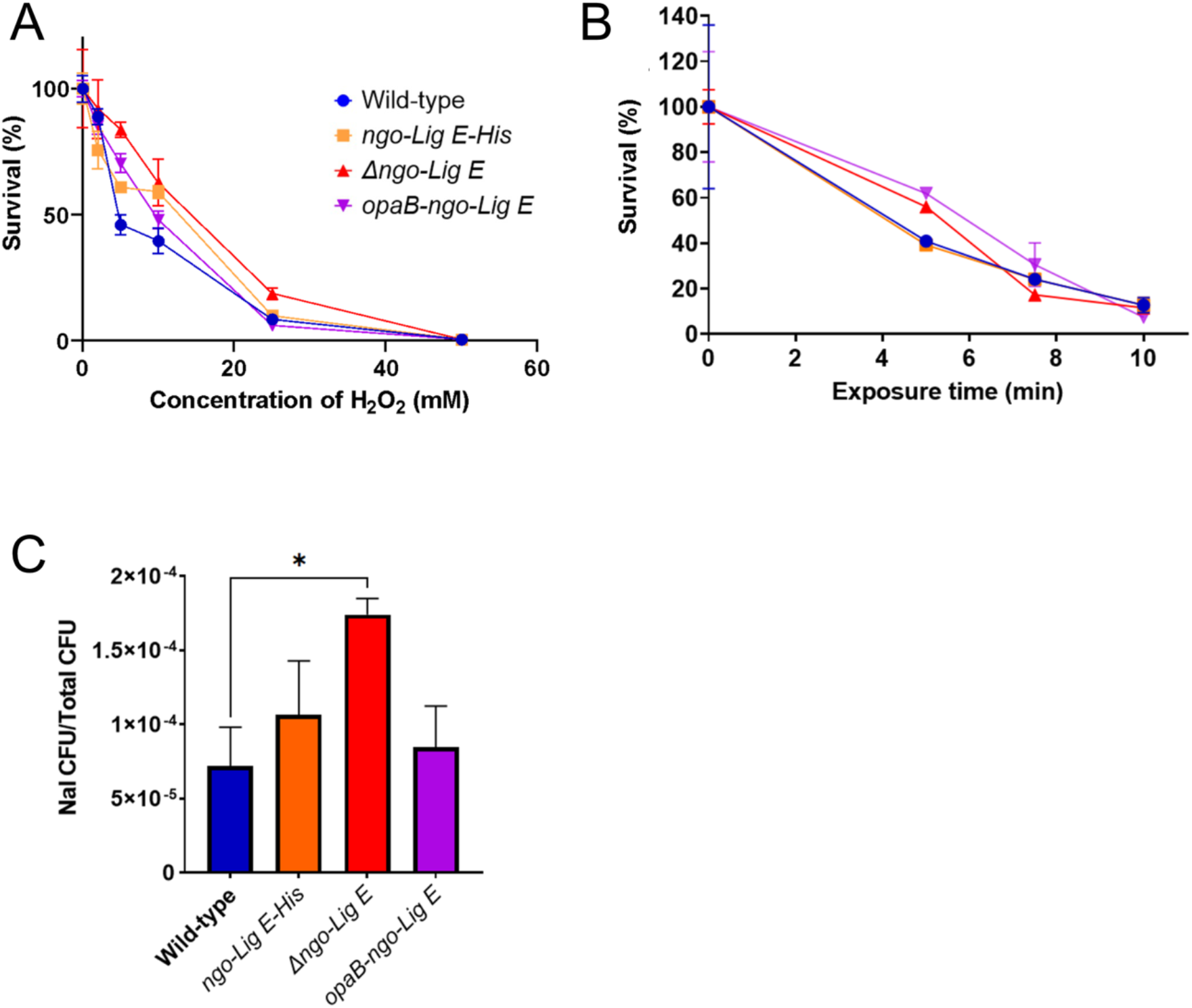
Damage and oxidative stress assays with N. gonorrhoeae strains. A) Survival after treatment with hydrogen peroxide. B) Survival after irradiation with UV light. C) Survival after treatment with nalidixic acid. Points are the mean of triplicate measurements and error bars represent the standard error of the mean. Significance values are given as * P ≤ 0.05. Comparisons which showed no significant difference (P >0.05) are not indicated.

### LIG E DELETION IMPACTS BIOFILM FORMATION AND CELL ADHESION

Due to its predicted periplasmic location, we considered whether Ngo-Lig E could influence biofilm formation. Crystal violet quantification of strains cultivated for 24 hours in 96 well plates indicated that deletion of Ngo-Lig E decreased the extent of biofilm produced by *Δngo-Lig E*; however, upregulating the expression of Ngo-Lig E did not increase biofilm production above that of the wild type (Figure 4 A). qPCR indicated that transcription of *ngo-lig E* was not upregulated in *wt* during biofilm formation relative to liquid culture which suggests that although deletion of Lig E diminishes biofilm it is not necessary to upregulate expression to produce biofilm (Figure 2).

**Figure 4.**
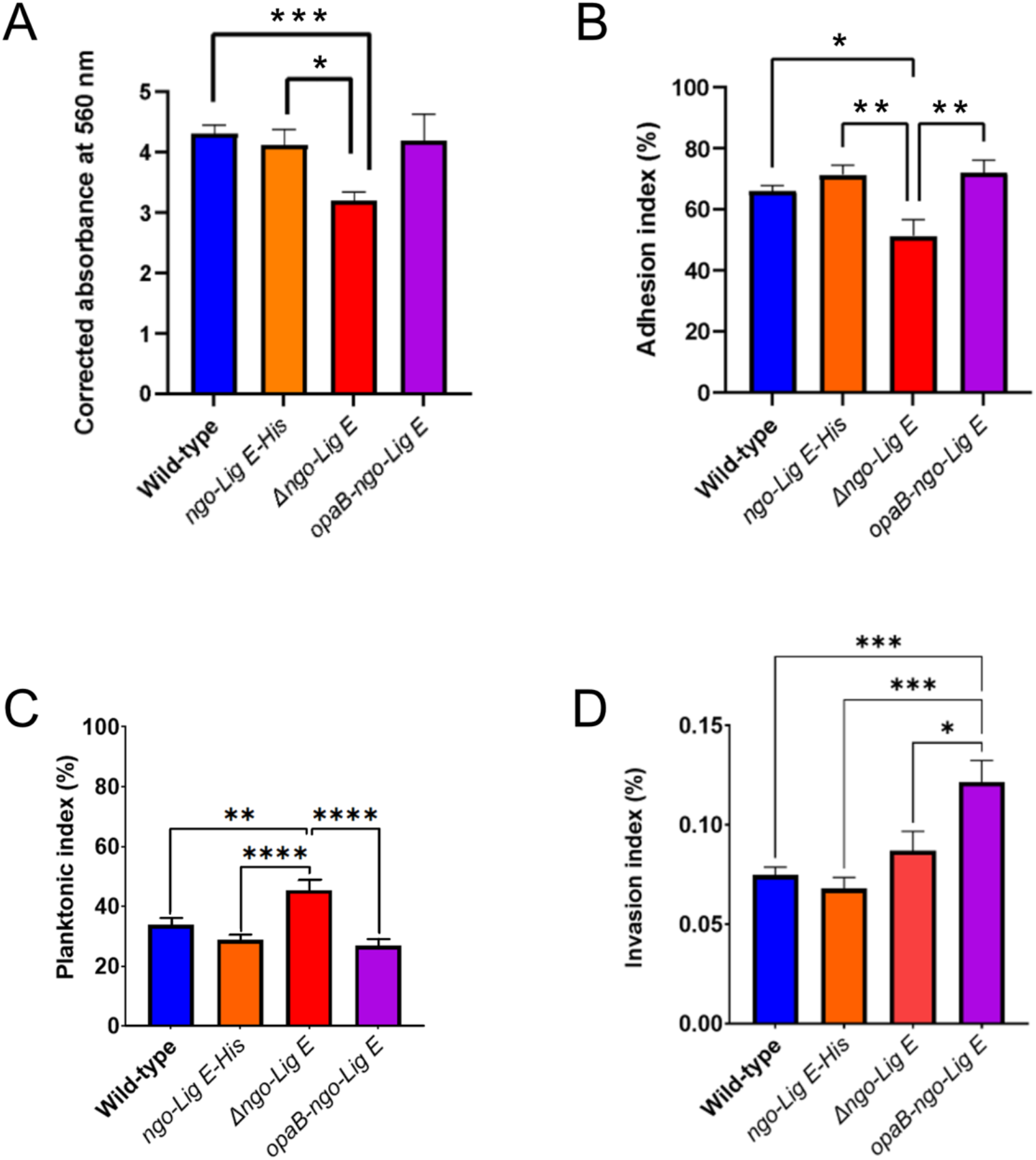
A) Biofilm formation in liquid culture by N. gonorrhoeae strains. B) Proportion of Ngo-Lig E mutants adhering to epithelial cells. C) Proportion of Ngo-Lig E mutants remaining in the non-adhered (planktonic) state when co-cultured with epithelial cells D) Proportion of Ngo-Lig E mutants invading epithelial cells relative to the total N. gonorrhoeae (planktonic and adhered). Points are the mean of triplicate measurements and error bars represent the standard error of the mean. Significance values are given as * P ≤ 0.05; ** P ≤ 0.01; *** P ≤ 0.001. Comparisons which showed no significant difference (P >0.05) are not indicated.

Given this result, we further asked whether the absence of Ngo-Lig E could influence the adhesion behaviour or infectivity of *N. gonorrhoeae* to human epithelial cells. Comparison of adhesion on endocervical cells indicates that *Δngo-Lig E* is impaired in its ability to attach to the eukaryotic cell surface, with a significantly greater portion of Ngo-Lig E knock-out cells remaining in the planktonic fraction (Figure 4 B and C); however, increased production of Ngo-Lig E did not enhance cell adhesion. Conversely, the decreased adhesion of the Ngo-Lig E deficient strain did not translate into defects in infectivity, however the strain over expressing Ngo-Lig E had increased rates of cell invasion (Figure 4 D). There were no significant differences in the behaviour of the His-tagged *ngo-Lig E-His* strain compared to the *wt* for cell infection, adhesion or biofilm formation.

### ATP-DEPENDENT NICK-SEALING ACTIVITY AND STRUCTURE OF NGO-LIG E

To compare the substrate specificity of Ngo-Lig E to that of other previously-characterised species, and to examine whether the His-tag impacts activity, mature Ngo-Lig E without the predicted N-terminal periplasmic leader sequence was recombinantly expressed and purified. *In vitro* assays with 5’ FAM labelled substrates demonstrated that Ngo-Lig E has highest activity on singly-nicked DNA. It is also able to join double-stranded breaks with 4 base-pair (bp) cohesive overhangs and has detectable activity on substrates with a mis-matched base-pair at the 3’OH end of the break (Figure 5 A and B). Maximal activity was observed at pH 7.1 and while significant joining was seen at higher pH, little to no activity was observed below this maximum (Figure 5 C). Inclusion of a C-terminal His-tag had no deleterious effect on Ngo-Lig activity with nicked DNA substrates, and actually appeared to enhance ligation efficiently (Supplement 7).

**Figure 5.**
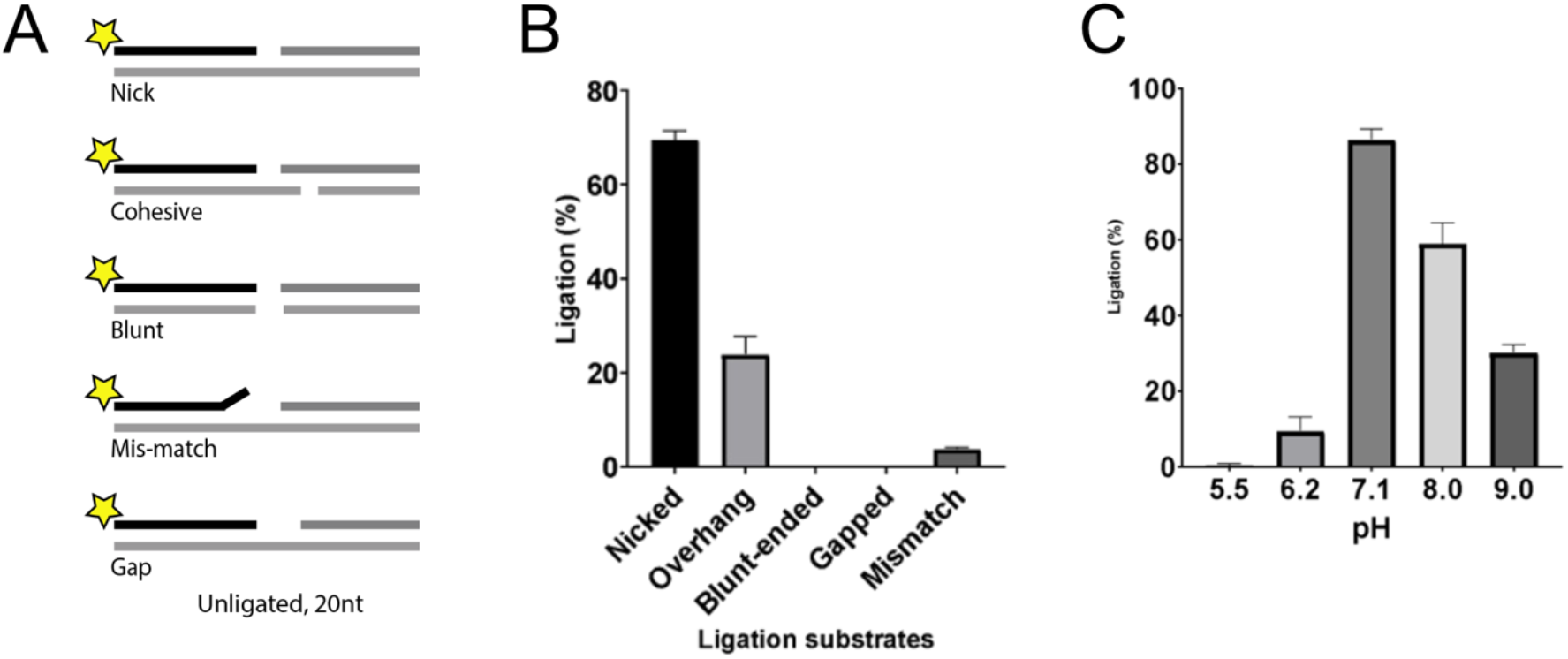
A) Schematic of FAM-labelled DNA substrates with ligatable breaks, B) Specific activity of Ngo-Lig E on different double-stranded breaks. C) pH dependence of Ngo-Lig E specific activity on singly-nicked DNA. Values represent the percentage of total substrate ligated quantified from band intensities and are the mean of three replicates. Error is the standard error of the mean.

To enable structural comparison with other Lig E proteins, Ngo-Lig E was crystalised in the presence of a 21 base-pair (bp) piece of DNA with a centrally placed phosphorylated nick. The resulting structure shows Ngo-Lig E in an open conformation bound to the DNA via interactions with its OB domain only (Figure 6 A). Covalent adenylation at Lys 22 is present in the NTase domain, indicating that we have captured a pre-step 2 state before complete encirclement of the DNA which would position the nick in the active site (Figure 6 B).

**Figure 6.**
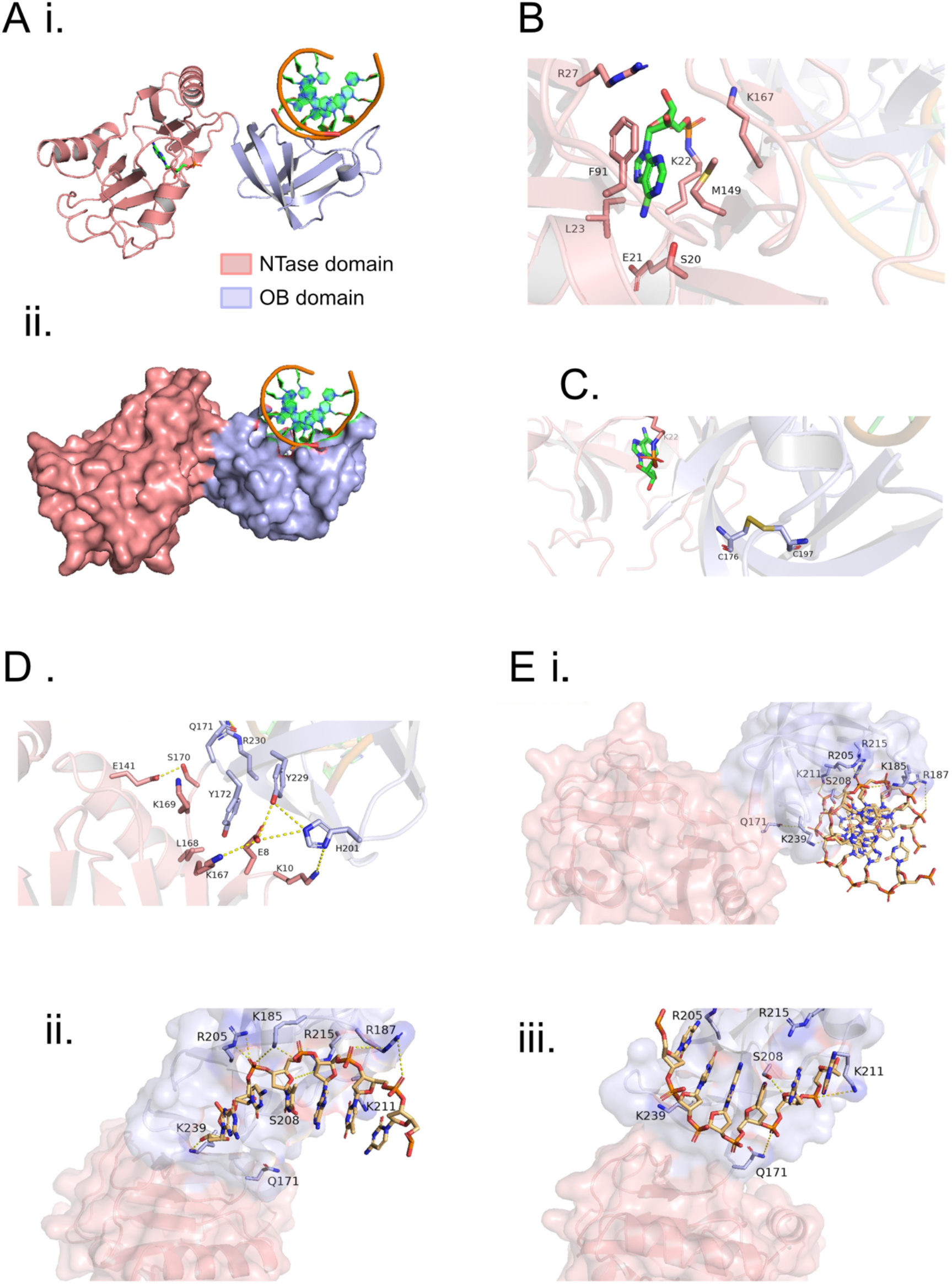
Structure of Ngo-Lig E. A) Overall structure of DNA-engaged Ngo-Lig E coloured by domain shown as a cartoon (i) or as a surface (ii). B) View of the NTase domain active site including the adenylated catalytic lysine residue. C) Position of the disulphide bond in the OB domain. D) inter-domain interactions surrounding the linker. Electrostatic and polar interactions less than 5Å are indicated by dashed yellow lines. E) Interaction of Ngo-Lig E with DNA duplex (i), specific interactions between conserved residues and the ‘complement’ strand of the duplex (ii), specific interactions between residues of the OB domain and the equivalent of the ‘nicked’ strand where the DNA break is located in other DNA-bound ligase structures (iii).

Analysis of the Ngo-Lig E structure confirms the presence of the disulphide bond between Cys 176 and Cys 197 in the OB domain which was previously predicted by computational modelling (Figure 6 C and Supplement 9 A)^6^. Examination of the linker region reveals a network of polar and electrostatic interactions that stabilise the open conformation of the domains including Lys 176 and Ser 170 from the linker with Glu 141 from the NTase domain; Tyr 172, Tyr 179 and His 201 from the OB domain with Lys 165, Glu 8 and Lys 10 of the NTase domain; Gln 171 of the linker with Arg 230 of the OB domain (Figure 6 D).

Despite the use of a 21-mer DNA fragment in the crystallization condition, only 6 base-pairs were visible in the final structure. Examination of crystal packing (Supplement 8) shows that the DNA forms a continuous filament throughout the crystal with protein subunits arranged along it. We suspect that, in the absence of the specific interactions imparted by nick-binding from the NTase domain, the Ngo-Lig E subunits have assembled non-specifically along the DNA filament; thus the 6 bp fragment in our structure represents a sample of the entire 21-mer piece. Key interactions with the DNA primarily involve basic residues from the OB domain and linker including Lys 239, Arg 205, Lys 185, Arg 215, Lys 211 and Arg 187.

These residues contact the backbone phosphates of the ‘complement’ strand which is base-paired opposite the nicked strand in other ligase-DNA structures (Figure 6 E ii.). There are fewer interactions with the ‘nick’ strand with the only contacts being Lys 239, Gln 171 and Ser 208 (Figure 6 E iii.). The electron density is continuous in this region of the phosphodiester backbone which is consistent with the ligase engaging an unbroken section of the DNA in a non-specific manner (Supplement 9 B). The nucleobases that are modelled in the structure represent the 6-mer combination giving the lowest R-free value during refinement and are in a region of the original 21-mer substrate before the nick; however, the density between base pairs is more symmetrically-distributed than would be expected for a well-ordered purine-pyrimidine pair, again supporting our suggestion that we have sampled an average of the 21-mer sequence in the present structure (Supplement 9 C). Comparison of the Ngo-Lig E structure with Lig E from *Alteromonas mediterranea* (Ame-Lig E) in a DNA-bound closed conformation shows the same conserved OB-domain residues are involved in DNA binding, despite the differences in overall conformation (Figure 7 A and B). Meanwhile superposition of Ngo-Lig E with Lig E from *Psychromonas sp*. SP041 (Psy-Lig E) which was crystalised in the open-state shows the domains of both proteins are in an identical configuration, despite the absence of DNA in the latter (Figure 7 C).

**Figure 7.**
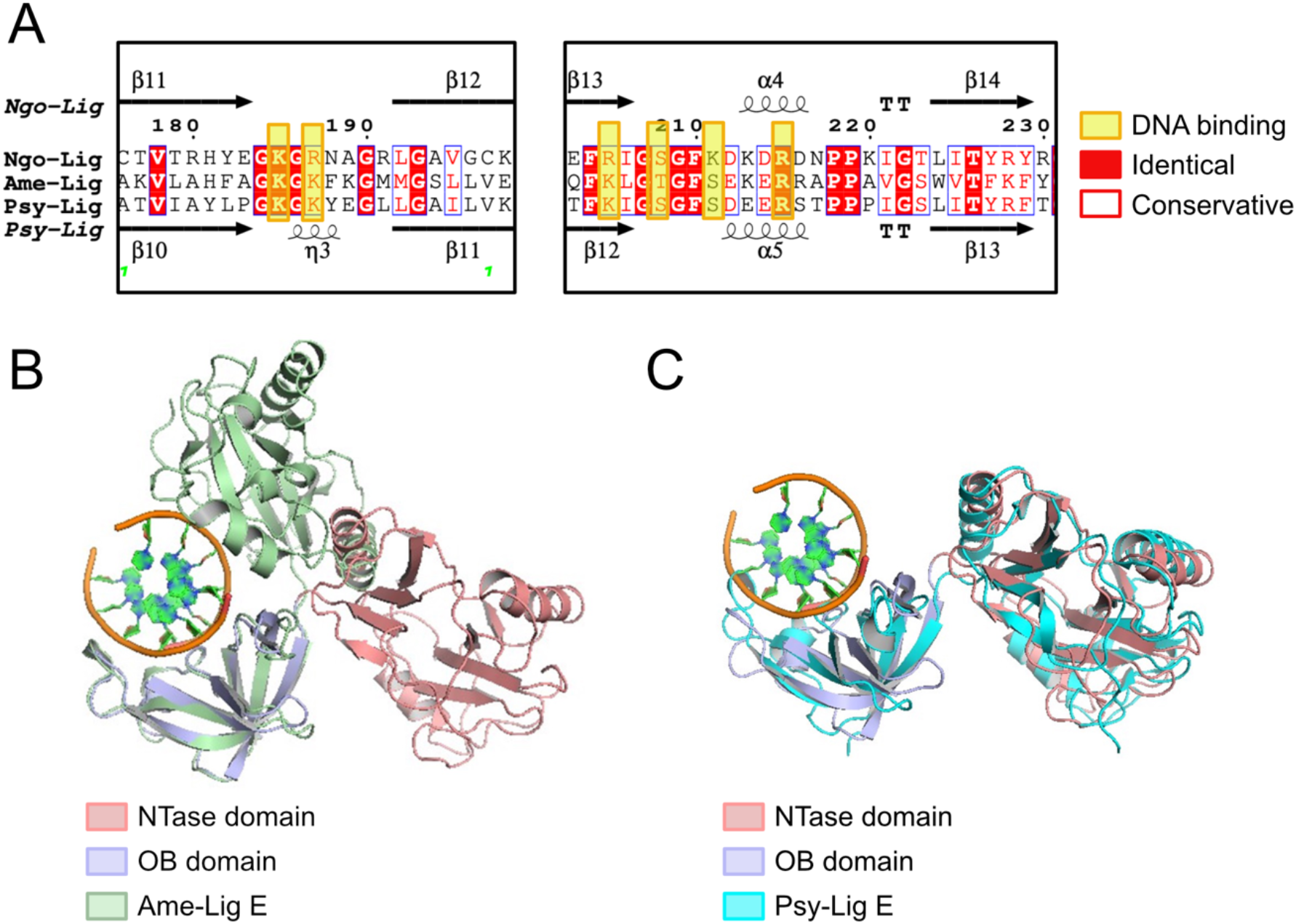
A) Alignment of other Lig E proteins that have been structurally characterised from Alteromonas mediterranea (Ame-Lig E) and Psychromonas sp. SP041 (Psy-Lig E). Structures were superimposed by alignment of the OB domains. DNA shown is from the Ngo-Lig E structure.

## DISCUSSION

Despite extensive *in vitro* characterisation of Lig E from a range of pathogenic and environmental Gram-negative bacteria ^7-11,37,38^ and the availability of structures of Lig E in both DNA-bound and DNA-free states ^9,10^ little is known about its biological function. Here we have examined the effect of knocking out Lig E in *N. gonorrhoeae*, as well as the impact of inserting a second copy of Lig-E under control of the strong constitutive Opa promoter.

Our results demonstrate that neither deletion nor upregulation of Ngo-Lig E impacts *N. gonorrhoeae* survival in liquid culture, or its planktonic growth rate. The dispensability of Ngo-Lig E under these conditions is counter to earlier studies which ascribed the inability to generate a viable deletion in *Haemophilus influenzae* to *lig E* being an essential gene ^37^.

Although the reason for this discrepancy is not clear, all bacterial genomes identified as possessing *lig E* also include the canonical NAD-dependent *lig A* DNA ligase responsible for chromosomal replication and most highly-conserved DNA repair pathways arguing for a non-essential function for *lig E* ^7^.

Similarly, Ngo-Lig E does not appear to mitigate the effects of oxidative stress and UV irradiation, both of which are anticipated to damage chromosomal DNA. Hydrogen peroxide treatment is a physiologically-relevant stressor as reactive oxygen species are produced during the host inflammatory response and by commensal *Neisseria* in the urogenital tract ^39^. Although UV exposure is not a natural source of damage in the urogenital mucous membranes that *N. gonorrhoeae* inhabits, this serves as a convenient proxy for generic DNA damage. The lack of its participation in DNA-damage survival is consistent with the predicted periplasmic location of Ngo-Lig E based on its N-terminal leader sequence, indicating that its primary substrate is not breaks or damages in chromosomal DNA. The enhanced survival of the *lig E* knock-out to treatment with nalidixic acid relative to the wild-type is somewhat counterintuitive; however, an equivalent but even starker example of this has been reported with knock-outs of the *phrB* ortholog which modulates DNA supercoiling ^40^. There, the striking 7,000-fold increased survival of nalidixic acid treatment was ascribed to mis-regulation of DNA topology by functional PhrB in the presence of DNA breaks caused by the treatment. In the case of *ngo-Lig E* deletion, we propose that improper recruitment of functional Ngo-Lig E to nalidixic acid-induced DNA breaks may interfere with resolution of this damage by canonical repair mechanisms. Previous *in vitro* studies have demonstrated that Lig E has a high affinity for nicked double-stranded DNA and we anticipate Ngo-Lig E will bind tightly to the doubly-nicked chromosomal DNA caused by nalidixic acid-poisoning of the gyrase B protein ^9^. The difference in the effect of nalidixic acid compared with UV and hydrogen peroxide may be due to their modes of action. UV and hydrogen peroxide typically impact the nitrogenous bases by oxidation or generation of dimerised photoproducts, but do not directly cause breaks in the phosphodiester backbone and therefore would not provide high-affinity binding sites for Lig E prior to initiation of repair pathways ^41^. Appending a 6-His tag to the C-terminus of native Ngo-Lig in the *N. gonorrhoeae* chromosome had essentially no impact on the growth or survival characteristics, suggesting that neither the tag nor the kanamycin selection cassette were deleterious to the strains. The slightly elevated levels of nick-sealing activity in the recombinant His-tagged enzyme relative to its untagged counterpart may be due to the increased positive charge in the C-terminal OB domain enhancing the enzyme’s interaction with the DNA substrate, or from minor differences in the purification protocol.

On the basis of Ngo-Lig E’s predicted periplasmic/ extracellular location, we investigated its impact on biofilm formation. Extracellular DNA is an important structural component of gonococcal biofilm and is derived from the *N. gonorrhoeae* chromosome either secreted through the T4SS system or released by autolysis ^16,19^. The role of the extracellular thermonuclease Nuc is well established in modulating the properties of gonococcal biofilm, with deletion of this gene resulting in thicker higher biomass biofilm as well as inability to escape chromatin-rich neutrophil extracellular traps (NETs) ^22,42^. Our results which indicate that Ngo-Lig E knock-outs produce less biofilm suggests that Ngo-Lig E acts in opposition to Nuc, potentially linking double-stranded segments of DNA. Although Ngo-Lig E is incapable of joining blunt-end double-stranded breaks, it demonstrated robust cohesive end joining as well as some activity on mismatched breaks, indicating it could potentially link regions of microhomology.

In addition to decreased biofilm production, our results indicate the Ngo-Lig E knock-out is less effective at adhering to epithelial cells, although it is not impaired in cell invasion. The cell adhesion process of *N. gonorrhoeae* involves microcolony attachment to the host cell surface via the Type IV pilus followed by interaction between *N. gonorrhoeae* opacity (Opa) proteins and epithelial receptors and other surface molecules ^43-47^. Although the rationale for Ngo-Lig E impacting cell adhesion is not clear, this could also be related to defects in biofilm formation as biofilm is suggested to play a role in cell-surface colonisation ^16,19^.

An emerging area of research in *N. gonorrhoeae* is the impact of biofilm architecture on dissemination of antibiotic resistance genes. In particular, the age and density of biofilm modulates the horizontal transfer of genes with more rapid spread observed in early biofilm and decreased dispersal in mature biofilms ^17^. Biofilm is also considered to provide a potential eDNA reservoir as the mobility of DNA fragments through this matrix is hindered by increasing DNA length and presence of DUS, leading to accumulation of *N. gonorrhoeae*-specific eDNA ^48^. In light of this, the potential of Lig E to modify the mobility of transformable eDNA by modulating biofilm properties, or potentially by acting on the transformation substrate itself by increasing its length or re-joining nicks resulting from nuclease or oxidative damage, is of considerable interest. One of the earliest studies of recombinant Lig E from *Neisseria meningitidis* suggested a possible function in competence for Lig E based on its periplasmic localisation signal and it has been noted that the majority of Lig E possessing bacteria are known to be competent in natural transformation and/or encode essential competence genes in their genomes ^7,38^. Immunological detection in the present study via an introduced His-tag was insufficient to define the extracellular or periplasmic distribution, however it is feasible that double-stranded DNA could serve as a substrate for periplasmically-localised Ngo-Lig E prior to import across the plasma membrane.

Finally, our DNA-engaged open-conformation structure of Ngo-Lig E reveals the specific interactions made between the OB domain and the DNA backbone, which are independent of sequence or phosphodiester-backbone continuity. Ngo-Lig E, like other members of this group of DNA ligases lacks any additional DNA binding domains or ‘latch’ regions such as those found in *Chlorella virus* type ligases which allow the enzyme to encircle the DNA duplex. Instead, Lig E ligases rely on the well-structured positively-charged surface of the OB domain for affinity. The conformation of the Ngo-Lig E enzyme-adenylate is consistent with the scanning mechanism previously proposed for locating breaks in the duplex ^49^. Here, non-specific interactions rapidly interrogate the substrate, and upon encountering a discontinuity in the duplex the ligase-adenylate re-orients its core domains about the linker region to position the active site for subsequent catalysis. The Ngo-Lig E structure suggests that in these minimal ligases, the OB domain is responsible for localising the enzyme to DNA and conformational scanning, prior to productive binding.

## CONCLUSIONS

Lig E is widely distributed among Beta-, Epsilon- and Gamma proteobacteria including some of the most prevalent human and agricultural pathogens; several of which are considered high-priority due to their emerging multi-antibiotic resistance. Our demonstration of a physiological role for Lig E in *N. gonorrhoeae* biofilm formation and cell adhesion recommends this enzyme for further study to understand its impact on virulence and pathogenicity, as well as potential roles in among commensal and environmental proteobacteria. Future directions will include more detailed phenotypic studies of the impact of Ngo-Lig E deletion on biofilm architecture and its interplay with other biofilm-modulating enzymes and processes. There is also the outstanding issue of the specific extracellular location of Ngo-Lig E, as well as exploration of a potential role in uptake and transformation of extracellular DNA which we anticipate will provide a more extensive picture of the biological function of this enigmatic DNA ligase protein.

## AUTHOR CONTRIBUTIONS

JP, JH and AW conceived of the study and experimental design, JP undertook construction and characterisation of N. gonorrhoeae mutants and recombinant Ngo-Lig E with guidance from JH and AW. JP and AW crystalised and solved the Ngo-Lig E structure. All authors participated in manuscript writing and approved the final version.

## DATA AVAILABILITY

Structure coordinates for the Ngo-Lig E protein-DNA complex have been deposited to the PDB with the identifier 8U6X.

## FUNDING

This work was supported by the Health Research Council of New Zealand (21/754 to AW; 19/602 and 23/534to JH) and Rutherford Discovery Fellowship (20-UOW-004 to AW). JP is supported by a University of Waikato Doctoral Scholarship.

## ACKNOWLEDGMENTS

This research was undertaken in part using the MX2 beamline at the Australian Synchrotron, part of ANSTO, and made use of the Australian Cancer Research Foundation (ACRF) detector.

## Notes

### Competing Interest Statement

The authors have declared no competing interest.

## REFERENCES

1 Płociński, P. et al. DNA Ligase C and Prim-PolC participate in base excision repair in mycobacteria. Nature communications 8, 1251 (2017). 10.1038/s41467-017-01365-y

2 Zhu, H. & Shuman, S. Gap filling activities of Pseudomonas DNA ligase D (LigD) polymerase and functional interactions of LigD with the DNA end-binding Ku protein. J Biol Chem 285, 4815–4825 (2010). 10.1074/jbc.M109.073874

3 Shuman, S. & Glickman, M. S. Bacterial DNA repair by non-homologous end joining. Nature Reviews Microbiology 5, 852–861 (2007). 10.1038/nrmicro1768

4 Ejaz, A., Goldgur, Y. & Shuman, S. Activity and structure of Pseudomonas putida MPE, a manganese-dependent single-strand DNA endonuclease encoded in a nucleic acid repair gene cluster. J Biol Chem 294, 7931–7941 (2019). 10.1074/jbc.RA119.008049

5 Ejaz, A. & Shuman, S. Characterization of Lhr-Core DNA helicase and manganese-dependent DNA nuclease components of a bacterial gene cluster encoding nucleic acid repair enzymes. J Biol Chem 293, 17491–17504 (2018). 10.1074/jbc.RA118.005296

6 Pan, J., Lian, K., Sarre, A., Leiros, H. S. & Williamson, A. Bacteriophage origin of some minimal ATP-dependent DNA ligases: a new structure from Burkholderia pseudomallei with striking similarity to Chlorella virus ligase. Scientific reports 11, 18693 (2021). 10.1038/s41598-021-98155-w

7 Williamson, A., Hjerde, E. & Kahlke, T. Analysis of the distribution and evolution of the ATP-dependent DNA ligases of bacteria delineates a distinct phylogenetic group ‘Lig E’. Mol Microbiol 99, 274–290 (2016). 10.1111/mmi.13229

8 Berg, K., Leiros, I. & Williamson, A. Temperature adaptation of DNA ligases from psychrophilic organisms. Extremophiles 23, 305–317 (2019). 10.1007/s00792-019-01082-y

9 Williamson, A., Grgic, M. & Leiros, H. S. DNA binding with a minimal scaffold: structure-function analysis of Lig E DNA ligases. Nucleic Acids Res 46, 8616–8629 (2018). 10.1093/nar/gky622

10 Williamson, A., Rothweiler, U. & Schroder Leiros, H.-K. Enzyme-adenylate structure of a bacterial ATP-dependent DNA ligase with a minimized DNA-binding surface. Acta Crystallographica Section D 70, 3043–3056 (2014). doi:10.1107/S1399004714021099

11 Williamson, A. & Pedersen, H. Recombinant expression and purification of an ATP-dependent DNA ligase from Aliivibrio salmonicida. Protein Expres Purif 97, 29–36 (2014). 10.1016/j.pep.2014.02.008

12 Cehovin, A. & Lewis, S. B. Mobile genetic elements in Neisseria gonorrhoeae: movement for change. Pathogens and disease 75 (2017). 10.1093/femspd/ftx071

13 Unemo, M. & Shafer, W. M. Antimicrobial resistance in Neisseria gonorrhoeae in the 21st century: past, evolution, and future. Clinical microbiology reviews 27, 587–613 (2014). 10.1128/cmr.00010-14

14 Martín-Sánchez, M. et al. Clinical presentation of asymptomatic and symptomatic heterosexual men who tested positive for urethral gonorrhoea at a sexual health clinic in Melbourne, Australia. BMC infectious diseases 20, 486 (2020). 10.1186/s12879-020-05197-y

15 Unemo, M. et al. Gonorrhoea. Nature Reviews Disease Primers 5, 79 (2019). 10.1038/s41572-019-0128-6

16 Falsetta, M. L. et al. The Composition and Metabolic Phenotype of Neisseria gonorrhoeae Biofilms. Frontiers in microbiology 2, 75 (2011). 10.3389/fmicb.2011.00075

17 Kouzel, N., Oldewurtel, E. R. & Maier, B. Gene Transfer Efficiency in Gonococcal Biofilms: Role of Biofilm Age, Architecture, and Pilin Antigenic Variation. J Bacteriol 197, 2422–2431 (2015). 10.1128/jb.00171-15

18 Steichen, C. T., Shao, J. Q., Ketterer, M. R. & Apicella, M. A. Gonococcal cervicitis: a role for biofilm in pathogenesis. The Journal of infectious diseases 198, 1856–1861 (2008). 10.1086/593336

19 1 Greiner, L. L. et al. Biofilm Formation by Neisseria gonorrhoeae. Infection and immunity 73, 1964–1970 (2005). 10.1128/iai.73.4.1964-1970.2005

20 Zweig, M. et al. Secreted single-stranded DNA is involved in the initial phase of biofilm formation by Neisseria gonorrhoeae. Environ Microbiol 16, 1040–1052 (2014). 10.1111/1462-2920.12291

21 Phillips, N. J. et al. Proteomic analysis of Neisseria gonorrhoeae biofilms shows shift to anaerobic respiration and changes in nutrient transport and outermembrane proteins. PLoS One 7, e38303 (2012). 10.1371/journal.pone.0038303

22 Steichen, C. T., Cho, C., Shao, J. Q. & Apicella, M. A. The Neisseria gonorrhoeae biofilm matrix contains DNA, and an endogenous nuclease controls its incorporation. Infection and immunity 79, 1504–1511 (2011). 10.1128/iai.01162-10

23 Dillard, J. P. Genetic Manipulation of Neisseria gonorrhoeae. Current protocols in microbiology Chapter 4, Unit4A.2 (2011). 10.1002/9780471729259.mc04a02s23

24 Callaghan, M. M. & Dillard, J. P. Transformation in Neisseria gonorrhoeae. Methods in molecular biology (Clifton, N.J.) 1997, 143–162 (2019). 10.1007/978-1-4939-9496-0_10

25 Schmittgen, T. D. & Livak, K. J. Analyzing real-time PCR data by the comparative CT method. Nature protocols 3, 1101–1108 (2008). 10.1038/nprot.2008.73

26 Petersen, T. N., Brunak, S., von Heijne, G. & Nielsen, H. SignalP 4.0: discriminating signal peptides from transmembrane regions. Nature methods 8, 785–786 (2011). 10.1038/nmeth.1701

27 Sharma, J. K. et al. Methods for competitive enrichment and evaluation of superior DNA ligases. Methods Enzymol 644, 209–225 (2020). 10.1016/bs.mie.2020.04.061

28 Schneider, C. A., Rasband, W. S. & Eliceiri, K. W. NIH Image to ImageJ: 25 years of image analysis. Nat Meth 9, 671–675 (2012).

29 Aragão, D. et al. MX2: a high-flux undulator microfocus beamline serving both the chemical and macromolecular crystallography communities at the Australian Synchrotron. Journal of synchrotron radiation 25, 885–891 (2018). 10.1107/s1600577518003120

30 Winn, M. D. et al. Overview of the CCP4 suite and current developments. Acta crystallographica. Section D, Biological crystallography 67, 235–242 (2011). 10.1107/s0907444910045749

31 Kabsch, W. XDS. Acta Crystallographica Section D: Biological Crystallography 66, 125–132 (2010). 10.1107/S0907444909047337

32 Jumper, J. et al. Highly accurate protein structure prediction with AlphaFold. Nature 596, 583–589 (2021). 10.1038/s41586-021-03819-2

33 Adams, P. D. et al. PHENIX: a comprehensive Python-based system for macromolecular structure solution. Acta crystallographica. Section D, Biological crystallography 66, 213–221 (2010). 10.1107/s0907444909052925

34 McCoy, A. J. et al. Phaser crystallographic software. J Appl Crystallogr 40, 658–674 (2007). 10.1107/s0021889807021206

35 Afonine, P. V. et al. Towards automated crystallographic structure refinement with phenix.refine. Acta crystallographica. Section D, Biological crystallography 68, 352–367 (2012). 10.1107/s0907444912001308

36 Emsley, P., Lohkamp, B., Scott, W. G. & Cowtan, K. Features and development of Coot. Acta crystallographica. Section D, Biological crystallography 66, 486–501 (2010). 10.1107/S0907444910007493

37 Cheng, C. H. & Shuman, S. Characterization of an ATP-dependent DNA ligase encoded by Haemophilus influenzae. Nucleic Acids Research 25, 1369–1374 (1997).

38 Magnet, S. & Blanchard, J. S. Mechanistic and kinetic study of the ATP-dependent DNA ligase of Neisseria meningitidis. Biochemistry-Us 43, 710–717 (2004). 10.1021/bi0355387

39 Seib, K. L. et al. Defenses against oxidative stress in Neisseria gonorrhoeae: a system tailored for a challenging environment. Microbiology and molecular biology reviews : MMBR 70, 344–361 (2006). 10.1128/mmbr.00044-05

40 Cahoon, L. A., Stohl, E. A. & Seifert, H. S. The Neisseria gonorrhoeae photolyase orthologue phrB is required for proper DNA supercoiling but does not function in photo-reactivation. Mol Microbiol 79, 729–742 (2011). 10.1111/j.1365-2958.2010.07481.x

41 Cadet, J. & Wagner, J. R. DNA base damage by reactive oxygen species, oxidizing agents, and UV radiation. Cold Spring Harb Perspect Biol 5 (2013). 10.1101/cshperspect.a012559

42 Juneau, R. A., Stevens, J. S., Apicella, M. A. & Criss, A. K. A thermonuclease of Neisseria gonorrhoeae enhances bacterial escape from killing by neutrophil extracellular traps. The Journal of infectious diseases 212, 316–324 (2015). 10.1093/infdis/jiv031

43 Higashi, D. L. et al. Dynamics of Neisseria gonorrhoeae attachment: microcolony development, cortical plaque formation, and cytoprotection. Infection and immunity 75, 4743–4753 (2007). 10.1128/iai.00687-07

44 Winther-Larsen, H. C. et al. Neisseria gonorrhoeae PilV, a type IV pilus-associated protein essential to human epithelial cell adherence. Proc Natl Acad Sci U S A 98, 15276–15281 (2001). 10.1073/pnas.261574998

45 Quillin, S. J. & Seifert, H. S. Neisseria gonorrhoeae host adaptation and pathogenesis. Nature reviews. Microbiology 16, 226–240 (2018). 10.1038/nrmicro.2017.169

46 LeVan, A. et al. Construction and characterization of a derivative of Neisseria gonorrhoeae strain MS11 devoid of all opa genes. J Bacteriol 194, 6468–6478 (2012). 10.1128/jb.00969-12

47 Walker, E., van Niekerk, S., Hanning, K., Kelton, W. & Hicks, J. Mechanisms of host manipulation by Neisseria gonorrhoeae. Frontiers in microbiology 14 (2023). 10.3389/fmicb.2023.1119834

48 Bender, N., Hennes, M. & Maier, B. Mobility of extracellular DNA within gonococcal colonies. Biofilm 4, 100078 (2022). 10.1016/j.bioflm.2022.100078

49 Bauer, R. J., Jurkiw, T. J., Evans, T. C., Jr. & Lohman, G. J. Rapid Time Scale Analysis of T4 DNA Ligase-DNA Binding. Biochemistry-Us 56, 1117–1129 (2017). 10.1021/acs.biochem.6b01261

