## Supplementary for "A role for the ATP-dependent DNA ligase Lig E of *Neisseria gonorrhoeae* in biofilm formation"

SUPPLEMENTARY INFORMATION

*Supplement 1. Primers for mutant confirmation and sequencing*

| Primer name | Sequence (5' to 3') | Purpose |
|---|---|---|
| Primer_1_KO_FD | CGGGGAGAATTTCGTAACGT | Lig E forward primer |
| Primer_2_Rev | CACTTTTTGCAGTGCGGCA | Lig E reverse primer |
| Primer_3_GFP_FD | AACTTTTCACTGGAGTTGTCCC A | Lig E-GFP forward primer |
| Primer_4_His_FD | CACCACCACCACCACCACT | Lig E-His forward primer |
| Primer_seq_Lig-E_Fd | CAAATCAGGGTGCAGTTTTGGC GGAA | Lig E forward sequencing primer |
| Primer_seq_Lig-E_Rev | GCCCCGCCCATCACGGGCAGG AGCA | Lig E reverse sequencing primer |
| NS10_Construct_Fd | GTCCGCAATGCGCCGAAT | NS10 forward primer |
| NS10_Construct_Rev | GCCTGTCCGCGTCTGAAA | NS10 reverse primer |
| NS10_External_Fd | GTAACGGTTTTCAATGCC | NS10 forward sequencing primer |
| NS10_External_Rev | GATATGCGCGGACATTAT | NS10 reverse sequencing primer |

*Supplement 2. qPCR primers and probes*

| Oligonucleotide name | Sequence (5' to 3') | Purpose |
|---|---|---|
| Lig E primer Fd | CGTATTGGGACGGAAAGCA | Lig E gene forward primer |
| Lig E primer Rev | AATCTGCTCGAACTGACCA C | Lig E gene reverse primer |
| Lig E probe | CAAAGGCTTTACCGCGCAG TTTCC | Lig E gene probe |
| 16s primerFd | CTGGGATAACACTGACGTT CAT | 16s gene forward primer |

| | | |
|---|---|---|
| 16s primer Rev | GCAATCAAGTTGCCCAACA G | 16s gene reverse primer |
| 16s probe | AGTCCACGCCCTAAACGAT GTCAA | 16s gene probe |

*Supplement 3. Oligonucleotide sequences for generating ligatable double-stranded DNA substrates. Combinations used for different ligatable breaks are given in Supplement 4*

| Oligonucleotide | Composition |
|---|---|
| L1 | 5'-(6-carboxyfluorescein) AGGCCATGGCTGATATCGCA-3' |
| L2 | 5'-(phosphate) TAGGCATTCGAGCTCCGTCG-3' |
| L3 | 5'- CGACGGAGCTCGAATGCCTATGCGATATCGGCCATGGCC T -3' |
| L6 | 5'-CGACGGAGCTCGAATGCCTA-3' |
| L7 | 5'-(phosphate) TGCGATATCAGCCATGGCCT-3' |
| L8 | 5'-(phosphate) ATATCAGCCATGGCCT-3' |
| L9 | 5'-CGACGGAGCTCGAATGCCTATGCG-3' |
| L10 | 5'- CGACGGAGCTCGAATGCCTACGCGATATCAGCCATGGCC T-3' |
| L11 | 5'- CGACGGAGCTCGAATGCCTAGTGCGATATCAGCCATGGC CT-3' |

*Supplement 4. Oligonucleotide combinations used to generate different double-stranded ligatable substrates. Sequences are given in Supplement 3.*

| Substrates | Ligatable strand | Complementary strand |
|---|---|---|
| Single nick | L1, L2 | L3 |
| Blunt ended | L1, L2 | L6, L7 |
| Overhang | L1, L2 | L8, L9 |
| Mismatch | L1, L2 | L10 |
| Gapped | L1, L2 | L11 |

*Supplement 5. Table of gene expression measured by qPCR and expressed as fold changes between mutants in biofilm and planktonic cultures. Data were collected in duplicate, and error represents the standard error of the mean*

| | Planktonic | Biofilm |
|---|---|---|
| Δngo-Lig E | ND | ND |

| | | |
|---|---|---|
| ngo-Lig E-His | 1.022 (± 0.061) | 1.081 (± 0.008) |
| opaB-ngo-Lig E | 88.62 (± 1.282) | 120.423 (± 22.244) |

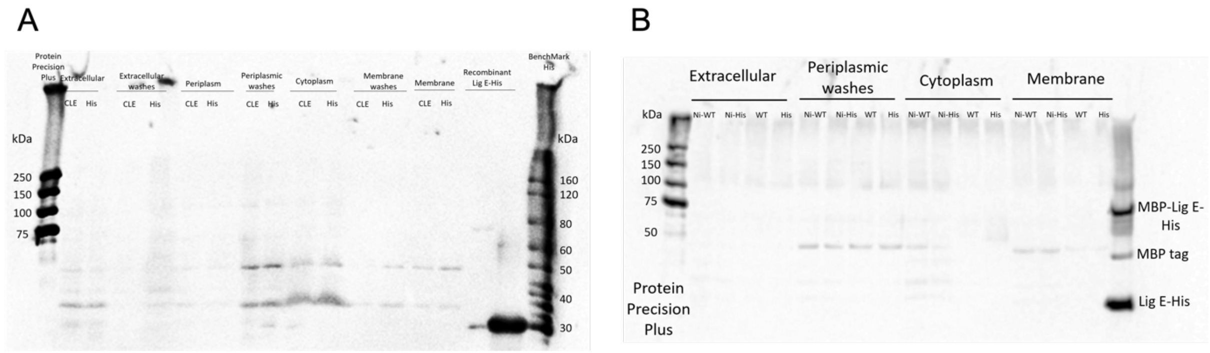


*Supplement 6. Western blots of different cellular fractions of N. gonorrhoeae MS11 using an anti-His antibody. Ladders are indicated in kDa. Sizes of the recombinant protein controls are 32kDa for Lig E-His, 44 kDa for the MBP tag and 76 kDa for MBP-Lig E-His. A) Comparison of the different fractions from the His-tagged constitutively upregulated Ngo-Lig E (opaB-Ngo-Lig E; CLE) and the His-tagged Ngo-lig E (ngo-Lig E-His; His) mutants. B) Comparisons of fractions from wild-type N. gonorrhoeae (WT) and the His-tagged Ngo-lig E (ngo-Lig E-His; His) as well as the respective nickel pull downs of each fraction (Ni-WT and Ni-His respectively).*

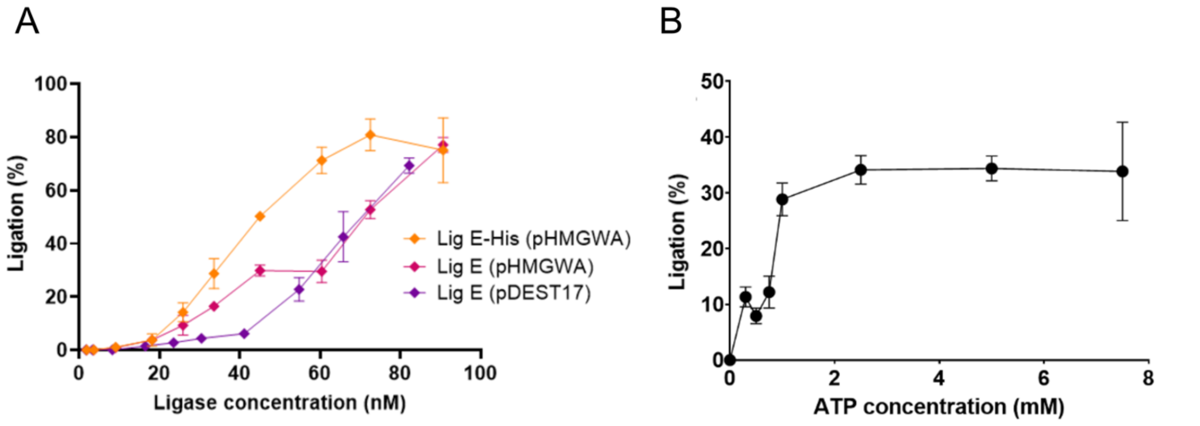


*Supplement 7. A) Ligase activity on singly-nicked DNA substrate for various recombinantly-produced Ngo-Lig E constructs; mature Ngo-Lig E purified from N-terminal His-tagged fusion (pDEST17) with the tag removed, Ngo-Lig E purified from N-terminal MBP-fusion (pHMGWA) with the tag removed and Ngo-Lig E purified from N-terminal MBP-fusion (pHMGWA) with the tag removed. B) ATP optimum of Ngo-Lig E. Measurements were made in triplicate, error represented the standard deviation of the mean.*

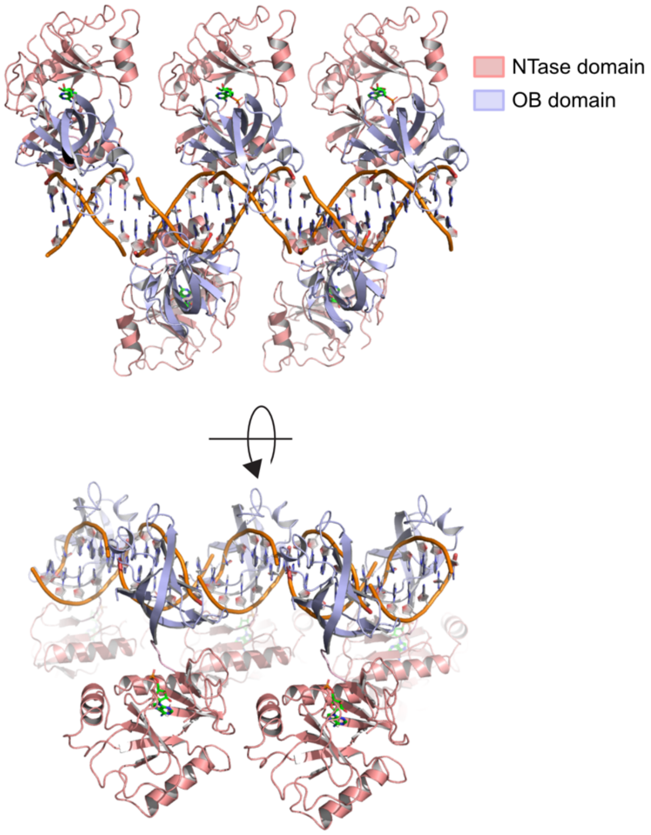


*Supplement 8. Symmetry-related molecules in the Ngo-Lig E crystal. Symmetry-related mates within 4 Å are shown generating the continuous DNA filament are shown.*

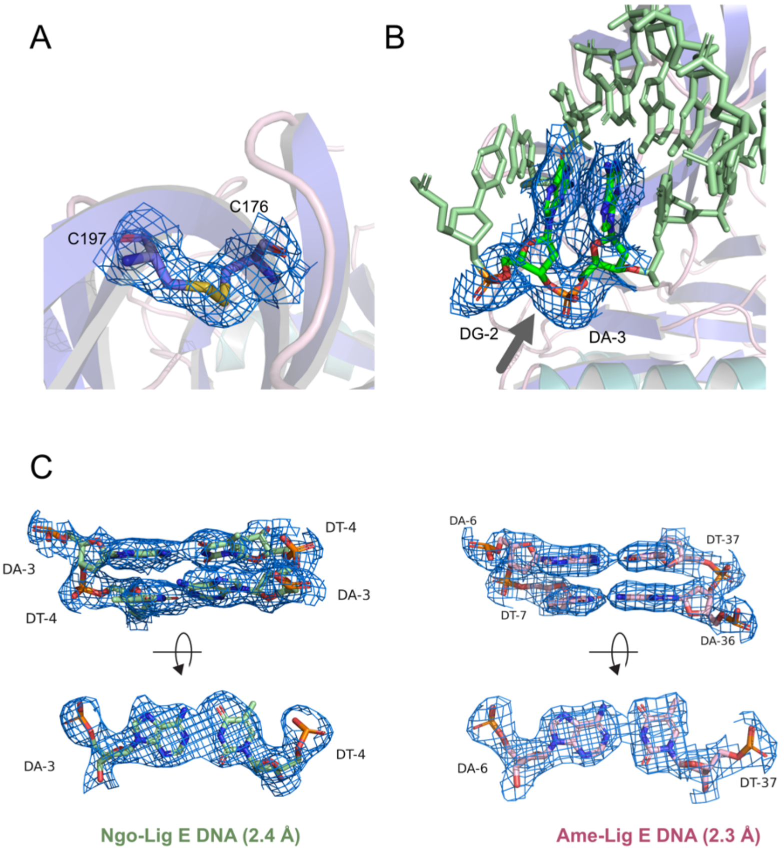


*Supplement 9. Observed electron density map (2Fo-Fc) surrounding key features of the Ngo-Lig E structure including: A) the disulphide bond, displayed at the 1.0 □ level. B) The DNA strand that is equivalent to the lick in other DNA ligase-DNA structures, shoing continuous density in the present structure. Map is displayed at the 0.5 □ level. C) Comparison of density surrounding centrally-placed base-pairs in the Ngo-Lig E structure and the equivalent base-pair in the Alteromonas mediterranea (Ame-Lig E) 6GDR.*
